# *Megasphaera* in the stool microbiota is negatively associated with diarrheal cryptosporidiosis

**DOI:** 10.1101/2020.10.01.323147

**Authors:** Maureen A. Carey, Gregory L. Medlock, Masud Alam, Mamun Kabir, Md Jashim Uddin, Uma Nayak, Jason Papin, A.S.G Faruque, Rashidul Haque, William A. Petri, Carol A. Gilchrist

**Author notes:** Corresponding Author: William A. Petri, Jr. 345 Crispell Ave, Carter Harrison Building Rm1709A, Charlottesville, VA 22908-1340. Alternate Corresponding Author: Rashidul Haque, Parasitology Laboratory, icddrb 68, Shaheed Tajuddin Ahmed Sharani, Mohakhali, Dhaka- 1212. Department of Pediatrics, University of Virginia, Charlottesville, USA.

## Abstract

**Background:** The protozoan parasites in the *Cryptosporidium* genus cause both acute diarrheal disease and subclinical (i.e. non-diarrheal) disease. It is unclear if the microbiota can influence the manifestation of diarrhea during a *Cryptosporidium* infection.

**Methods:** To characterize the role of the gut microbiota in diarrheal cryptosporidiosis, the microbiome composition of both diarrheal and surveillance *Cryptosporidium*-positive fecal samples was evaluated using 16S rRNA gene sequencing. Additionally, the microbiome composition prior to infection was examined to test whether a preexisting microbiome profile could influence the *Cryptosporidium* infection phenotype.

**Results:** Fecal microbiome composition was associated with diarrheal symptoms at two timepoints. *Megasphaera* was significantly less abundant in diarrheal samples when compared to subclinical samples at the time of *Cryptosporidium* detection (log_2_(fold change) = -4.3, *p*=10^−10^) and prior to infection (log_2_(fold change) = -2.0, *p*=10^−4^). Random forest classification also identified *Megasphaera* abundance in the pre- and post-exposure microbiota.as predictive of a subclinical infection.

**Conclusions:** Microbiome composition broadly, and specifically low *Megasphaera* abundance, was associated with diarrheal symptoms prior to and at the time of *Cryptosporidium* detection. This observation suggests that the gut microenvironment may play a role in determining the severity of a *Cryptosporidium* infection.

**Summary:** *Megasphaera* abundance in the stool of Bangladeshi infants is associated with the development of diarrhea upon infection with the *Cryptosporidium* parasite.

## INTRODUCTION

Protozoan parasites in the *Cryptosporidium* genus cause both acute diarrhea and subclinical (*i.e*. non-diarrheal) disease, and both clinical outcomes are associated with poor physical and neurocognitive growth in infants [1–6]. These parasites are the fifth leading cause of diarrhea in young children [7] and recent studies have estimated the global burden of *Cryptosporidium* diarrhea mortality to be as high as 50,000 deaths annually [8]. This burden is disproportionately borne by young children [9]. Importantly, no therapies exist to treat *Cryptosporidium* infection in children or immunocompromised individuals [10]. Thus, there is a pressing need to prevent cryptosporidiosis mortality.

Understanding the difference in the host, parasite, and environment during acute diarrheal and subclinical infections may reveal new therapeutic solutions. Human polymorphisms are associated with an increased host susceptibility to cryptosporidiosis; however, these mutations do not completely explain the differences in infection outcomes [19,20]. Parasite genetics (within and across species) have been associated with differences in their host range [16,21–23]. The role of the microbiome upon infection by *Cryptosporidium* has been examined in healthy adults [24] and animals [25,26]; but its role in differentiating diarrheal and subclinical infections is not known and nor is the impact of any differences in the microbiome composition occurring during infant cryptosporidiosis.

Here, we interrogate the association between diarrheal status during cryptosporidiosis and a child’s microbiome using fecal samples from infants living in Mirpur and Mirzapur, Bangladesh. In Mirpur, *Cryptosporidium* diarrhea was frequent (24% of infections) and detected *Cryptosporidium* species included *Cryptosporidium hominis, parvum*, and *meleagridis* with *C. hominis* as the most common. In contrast, most infections in Mirzapur were subclinical (98%) and *Cryptosporidium meleagridis* was the most common detected species [1]. Because *Cryptosporidium*-associated diarrhea was infrequent in Mirzapur and most infections involved *C. meleagridis* rather than *C. hominis* or *C. parvum*, the association between diarrheal status and microbiome composition in infants in Mirzapur could not be decoupled from an alternative infection phenotype caused by *C. meleagridis*. We therefore focused our analysis on Mirpur due to the variation in diarrheal status and the dominance of the *Cryptosporidium hominis* species in this population. We found that the microbiota demonstrated high variability between children but despite this, microbiota composition and a low abundance of *Megasphaera* were associated with diarrheal symptoms both at the time of *Cryptosporidium* detection and prior to infection. Thus, we propose that *Megasphaera* may prevent acute diarrhea during parasite infection or is a biomarker for other unknown protective factors.

## METHODS

### Cohort

Children were enrolled into a community-based prospective cohort study of enteric infections which was established at the urban and rural Bangladesh sites, Mirpur and Mirzapur respectively (**Figure 1A**, “Cryptosporidiosis and Enteropathogens in Bangladesh”; ClinicalTrials.gov identifier NCT02764918) [1,28]. Stool samples were collected monthly and during diarrheal episodes. Diarrhea was defined as three or more loose stools within 24 hours, as reported by the child’s caregiver. Both a pan-species and species-specific qPCRs were used to identify the species of the *Cryptosporidium* infecting the children (Steiner et al. 2018). If positive samples were collected with an interval of less than or equal to 65 days they were regarded as derived from one infection event [1,28]. In addition to the collection of stool samples, a study database was created containing clinical information on each episode of diarrhea a child experienced, antibiotic consumption, and anthropometric measurements as well as data on the household demographics [1]. A subset of the *Cryptosporidium*-positive and corresponding ‘pre-detection’ *Cryptosporidium*-negative surveillance samples were analyzed. The data from Mirzapur (**Figure 1A** *right*) were only included in the post-hoc analysis represented in Figure 4F due to the limited amount of information on the antibiotic history of these children, the rarity of diarrheal cases at this site, and the high prevalence of *C. meleagridis* at the site relative to the more common *C. hominis* species detected in Mirpur.

**Figure 1:**
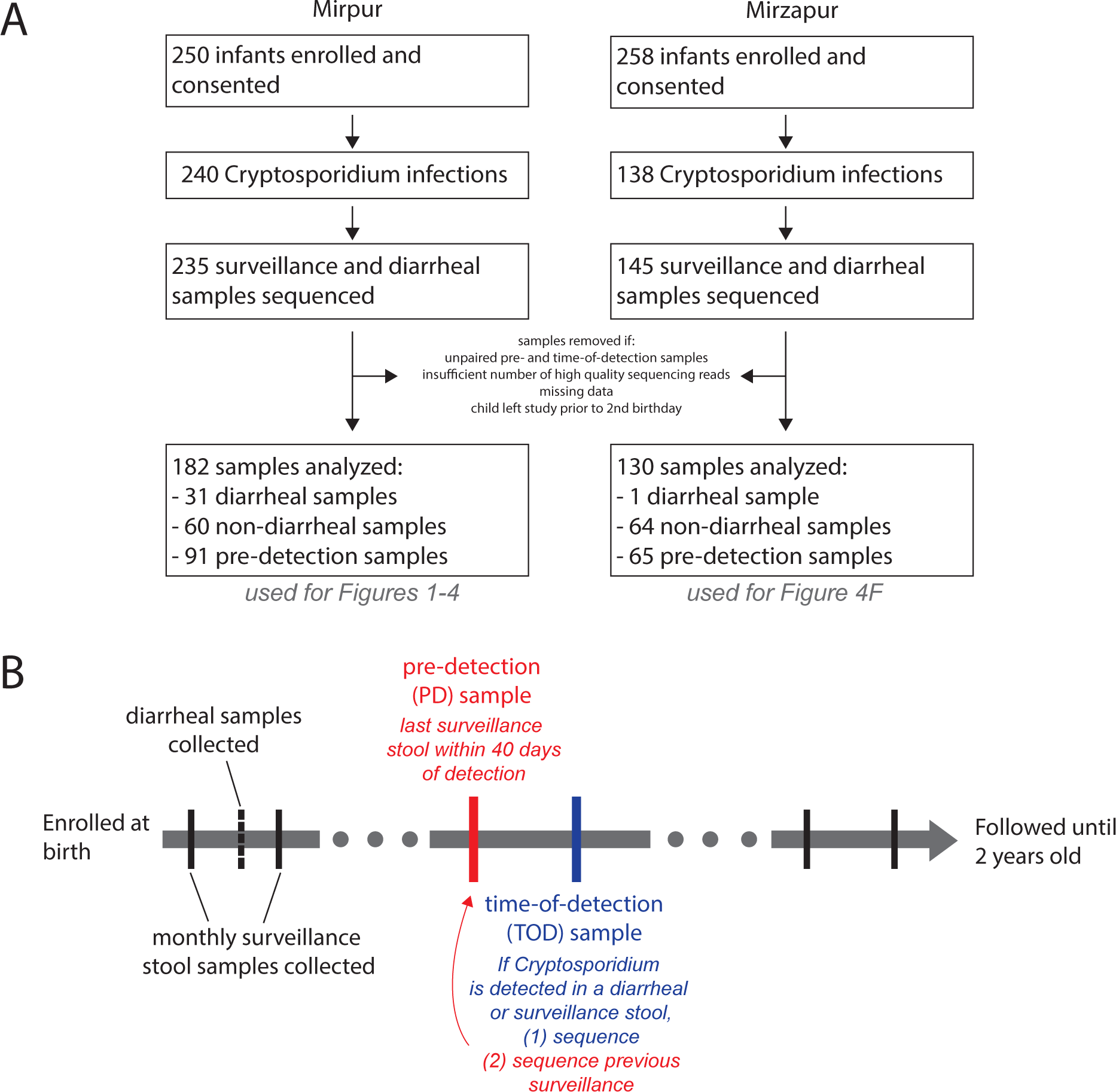
Study design. **A**: Overall cohort design and sample collection. For more information, see [1,28]. Samples from Mirzapur were only used in post-hoc analysis in **Figure 4F. B**: Paired samples were selected to assess *Cryptosporidium*-positive samples (time-of-detection, TOD) and the preceding surveillance sample (pre-detection, PD). *Cryptosporidium*-positive samples were identified from both monthly surveillance and diarrheal stool samples, generating our subclinical and diarrheal sample groups.

**Figure 4:**
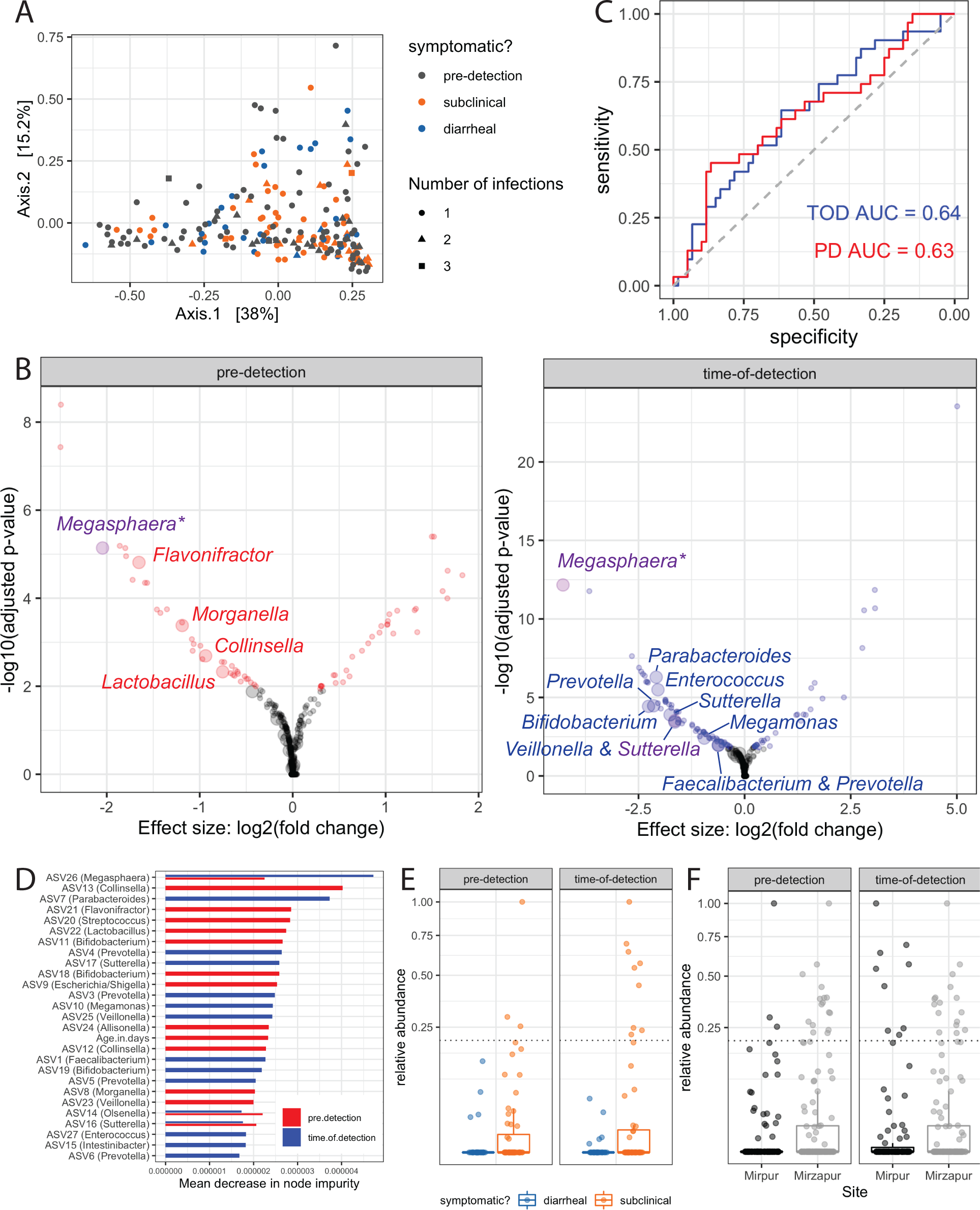
Identifying associations between diarrheal symptoms and the microbiota. **A**: Pre-detection and time-of-detection sample microbiota were indistinguishable via principal coordinate analysis using a permutational multivariate analysis of variance using distance matrices and a significance cutoff of p < 0.05, as were subclinical and diarrheal *Cryptosporidium*-positive samples. Principal coordinate analysis of ASV quantification across all samples using Euclidean distance. **B**: Univariate statistics identifies ASVs associated with symptoms in the pre-detection samples and time-of-detection samples. Statistically significant differential expressed ASVs are colored, whereas grey points represent ASVs that were not different or not significantly different, using DESeq2. Large points indicate ASVs that were also identified as important using random forest classification, whereas small points were not among the top 15 most important variables. Random forest classifiers were built to predict the presence of diarrhea upon *Cryptosporidium* infection. Importantly, purple points represent statistically significant ASVs that were also among the most important variables for classifiers made at both timepoints. **C**: Random forest classifiers were built from the time-of-detection (TOD) microbiota (*blue*) or pre-detection microbiota (*red*). Area-under-the-curve (AUC), a metric of classifier accuracy, is listed for each classifier. **D:** Most important variables, as ranked by mean decrease in node impurity (or, Gini importance), from the pre-detection and time-of-detection classifiers. Important variables were similarly important, within and across models. Of note, age was not an important variable in the time-of-detection classifier. **E:** One ASV assigned to the *Megasphaera* genus was significantly less abundant in diarrheal cases via univariate analyses (at both timepoints) and was among the top 15 most important variables for the classifiers for both timepoints. Relative abundance of each ASV is plotted for each sample with each box representing the median (inner line), 25^th^ percentile and 75^th^ percentile. Upper whiskers extend from the top of the box to the largest value within 1.5 times the interquartile range (distance between 25^th^ and 75^th^ percentile), and the lower whisker extends to the smallest value within 1.5 time the interquartile range. **F**: The *Megasphaera* ASV was also more likely to be high-abundance (above dashed line) in samples at the second study site, Mirzapur, where diarrheal cryptosporidiosis was less common when compared to Mirpur; however, environmental factors, including the causal *Cryptosporidium* species, were also different in Mirzapur [1]. Increased *Megasphaera* abundance in Mirzapur may partially explain reduced diarrhea associated with cryptosporidiosis in that community. ASVs: Amplicon sequence variants.

**Supplemental Figure 3:**
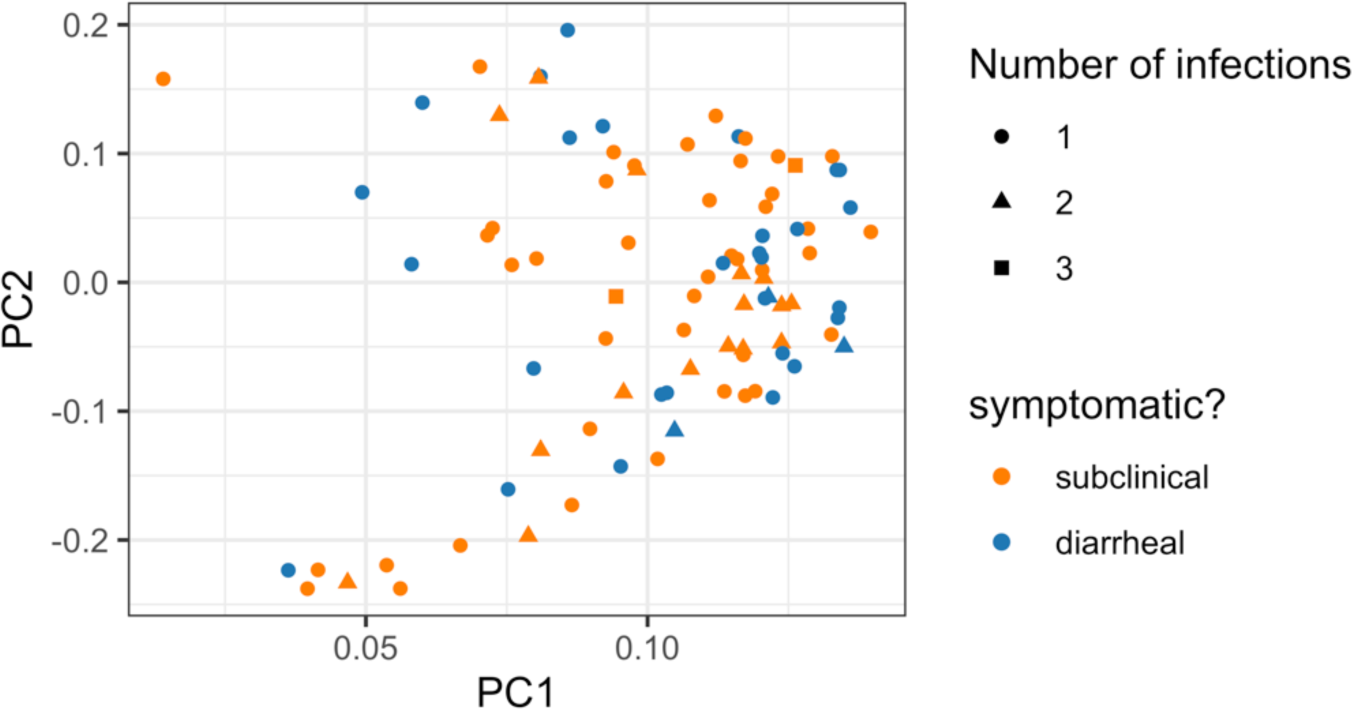
The change in ASV quantification (between pre-detection and time-of-detection samples) for subclinical and diarrheal samples were also indistinguishable via principal coordinate analysis. Principal coordinate analysis of the change in ASV quantification across all samples using Euclidean distance. Pre-detection and time-of-detection samples were run on different sequencing runs (see *Methods*) so the biological shift in microbiota might be confounded by the technical shift in run-to-run variation.

The study was approved by both the Ethical and Research Review Committees of the International Centre for Diarrhoeal Disease Research, Bangladesh, and by the Institutional Review Board of the University of Virginia. For each child, informed written consent was obtained from their parent or guardian.

### DNA extraction

On the day of collection, stool samples were brought to the study clinic and transported to our laboratory at 4°C, where they were aliquoted in DNase- and Rnase-free cryovials for storage at -80°C. For DNA extraction, samples were thawed and 200mg removed for total nucleic acid extraction; see [1]. To verify the extraction protocol, phocine herpesvirus (EVAg European Virus Archive Global, Erasmus MC, Department of Virology, Rotterdam, The Netherlands) and bacteriophage MS2 (ATCC 15597B; American Type Culture Collection, Manassas, VA) were added into each sample as positive controls.

### 16S ribosomal sequencing and processing

The V4 region of the 16S rRNA gene was amplified using the previously described phased Illumina-eubacteria primers and protocol from [29,30] with the minor modification that the illumina MiSeq v3 chemistry was used to generate 300bp paired-end reads. Sequencing was performed by the University of Virginia’s Genome Analysis and Technology Core. Negative controls included extraction blanks throughout the amplification and sequencing process. As positive controls, DNA was extracted from the HM-782D Mock Bacteria Community (ATCC through BEI Resources) and analyzed on each sequencing run (**Supplemental Figure 1A-C**). Additionally, a PhiX DNA library was added at 20% into each sequencing run to increase genetic diversity prior to parallel sequencing in both forward and reverse directions using the Miseq v3 kit and machine (per the manufacturer’s protocol).

**Supplemental Figure 1:**
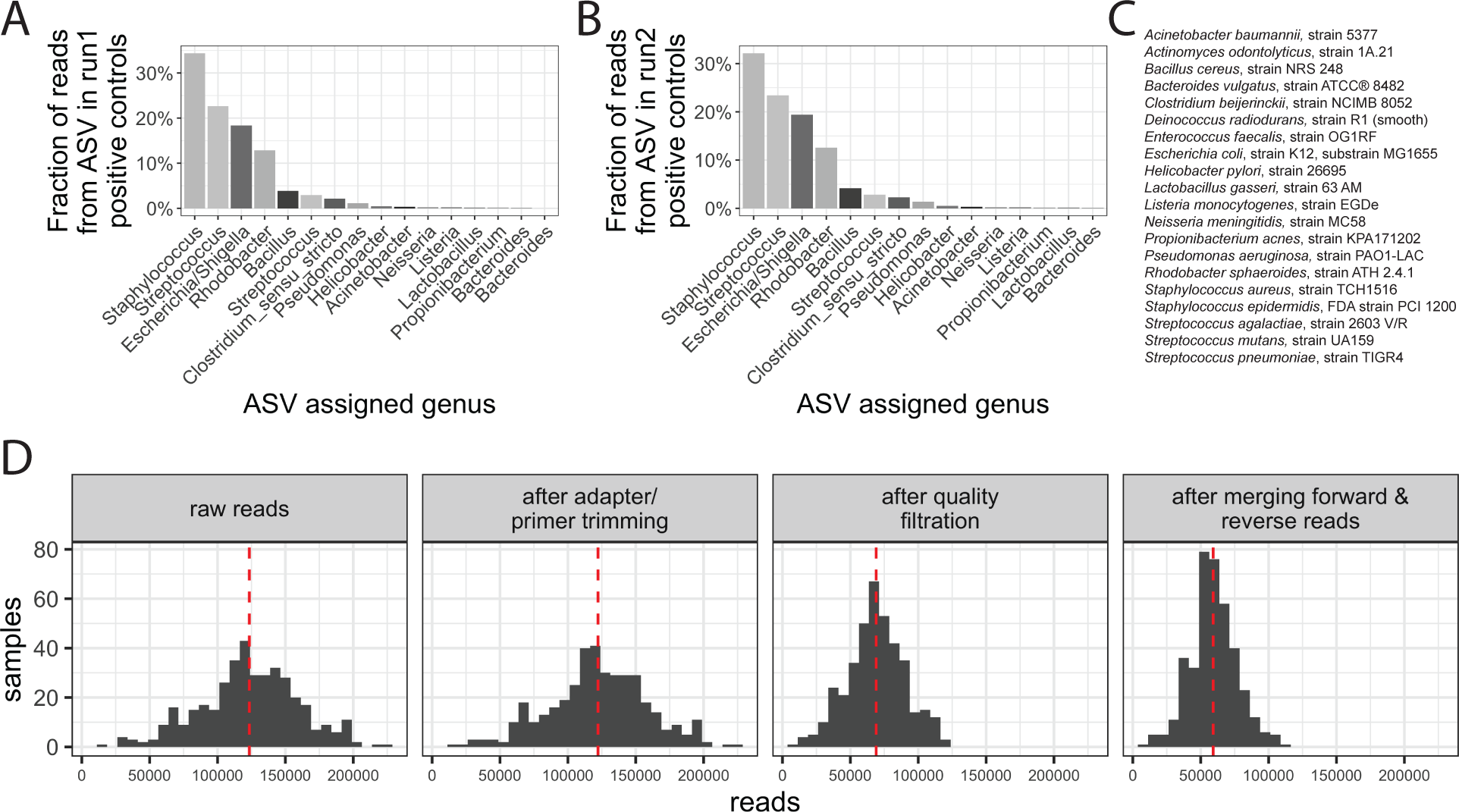
Sequencing controls and preprocessing. **A-B**: ASVs detected from run 1 (**A**) and run 2 (**B**) positive controls containing purified DNA from BEI Resources’ HM-782D mock community. **C**: Members of the HM-782D mock community. Positive controls for each run had the expected read assignments. **D:** Histogram of reads throughout the preprocessing pipeline; dotted line represents the distribution mean. Samples were sequenced deeply, permitting subsampling of 10,000 reads to remove biases introduced by variable read depth.

Sequencing produced 48,146,401 reads with a mean of 118,295.8 and median of 121,519 reads per sample (*raw reads* from **Supplemental Figure 1D**). Sequence adaptors were then removed using Bbtools [31] and primers were removed using CutAdapt [32]; quality-based filtering was performed with DADA2 [33]. Quality filtration reduced the total number of reads to a mean of 59,202.2 reads per sample (**Supplemental Figure 1D**). In brief, reads were removed and trimmed based on overall read quality and base pair quality: forward and reverse reads were trimmed to 250 or 200 base pairs and removed if there were more than 3 or 6 expected errors, respectively. Reads were also truncated at the first instance of a quality score (Phred or Q score) of 2 or less. Next, forward and reverse reads were merged with only 1 mismatch permitted. Lastly, taxa assignments were made using DADA2’s naive Bayesian classifier method and the Ribosomal Database Project’s Training Set 16 (release 11.5) reference database [33] and reads that did not map to bacteria (including human contaminants, archaea, and mitochondrial or chloroplast DNA) were removed, resulting in a mean of 27,809 reads per sample.

Samples with fewer than 10,000 reads and unpaired samples (those with no pre-detection or time-of-detection sample within 42 days) were removed from consideration; all were subsampled to a uniform depth of 10,000 reads per sample to correct for differences in sequencing depth across samples. Following these filtration and processing steps, 2953 amplicon sequence variants (ASVs) and 182 stool samples remained in the dataset.

The 182 paired pre-detection and time-of-detection samples (91 pre- and 91 post-detection), as well as additional positive and negative control samples (amplification blanks) and additional samples that did not pass our selection criteria, were split into two sequencing runs to increase the sequencing depth. The first sequencing run included all pre-detection samples and the second sequencing run included all time-of-detection samples. As an unintentional result of this choice, sequencing batch effects may result in spurious differences between pre- and time-of-detection samples, thus analyses are focused on symptomatic vs. subclinical samples within each time point (i.e., within the same sequencing batch).

### Statistical and machine learning analyses

All of the following data processing and statistical analyses were performed in R [33–37]; see **Supplemental Material** for code and software versions. Appropriate statistical tests were selected and are described as introduced throughout the Results.

For machine learning analyses, random forest analysis was used to classify subclinical or diarrheal samples using associated metadata and/or amplicon sequence variant (ASV) abundances, and the trained classifiers were used to identify individual variables that were important for prediction accuracy [38]. Within a random forest classifier, individual trees were built from subsets of the data and model performance was evaluated by predicting the class of each sample using only the trees in the random forest that were not constructed using that sample (i.e., out-of-bag performance). Here, variables were ranked by their effect on classifier certainty, which influenced overall accuracy, using the mean decrease in node impurity (via the Gini coefficient). Variables that maximally split samples by classification group yielded a larger forest-wide node impurity and, thus, more important variables had a higher mean decrease in node impurity. Analytic code is provided in the supplemental material; analyses and figure generation were performed in R [37,39–51].

## RESULTS

### Prevalence of diarrhea and antibiotic use

Infants were enrolled into a prospective cohort from Mirpur, Dhaka, Bangladesh to study enteric infections (**Figure 1A**, *left*); this cohort was part of a larger assessment of diarrhea in Bangladesh, published previously (**Figure 1A;** [1]). Each child was monitored by community health workers for enteric disease, including collection of monthly surveillance and diarrheal stool samples during the first two years of life. Diarrhea and antibiotic use were common in this cohort (**Figure 2A** & **Supplemental Figure 2**), and *Cryptosporidium* spp., including *C. hominis* and *meleagridis*, were frequently detected during diarrhea (**Table 1**). These parasites cause both subclinical and overt diarrheal infections [1].

**Table 1:**
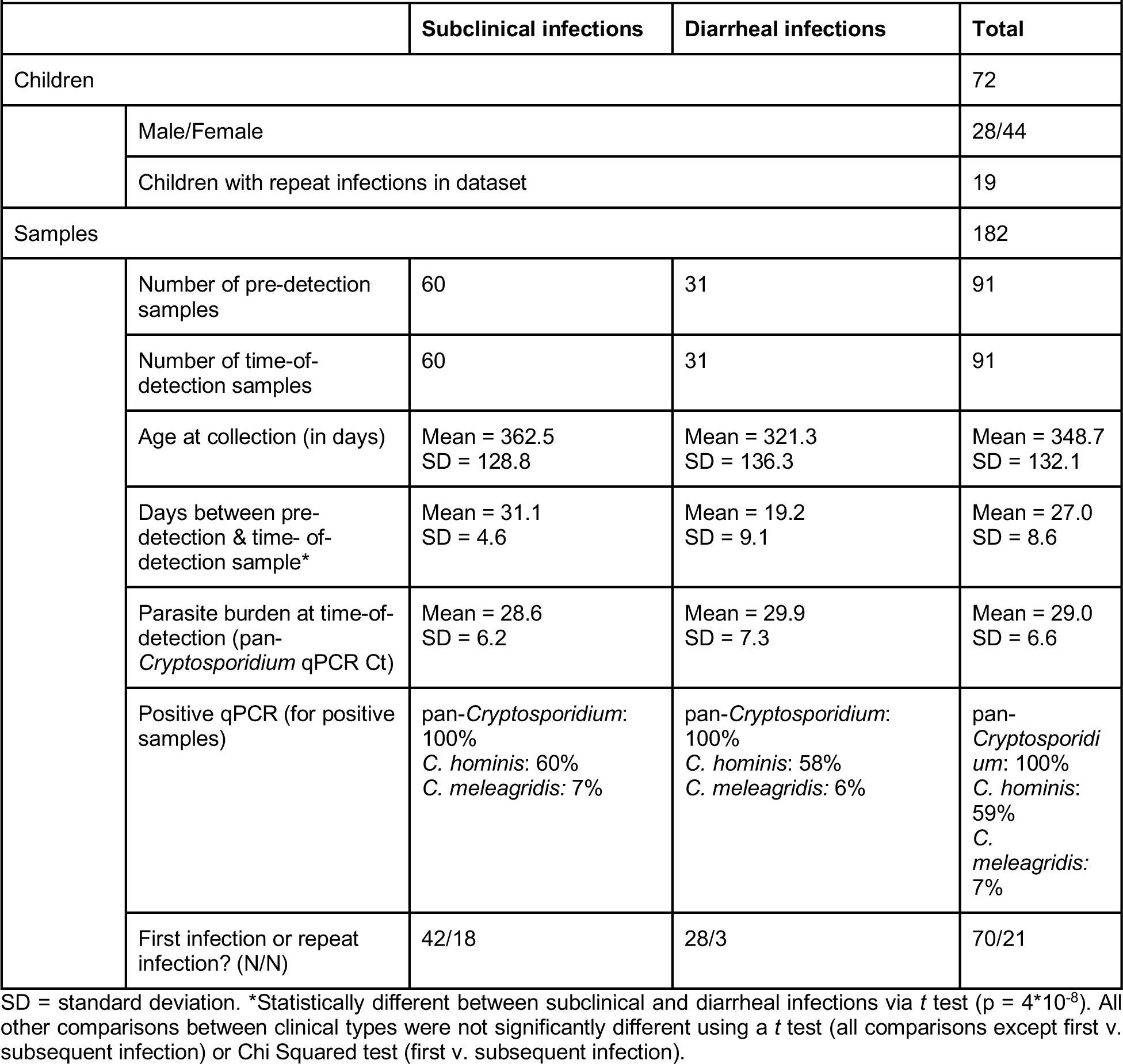
Sample summary statistics for samples from Mirpur.

**Figure 2:**
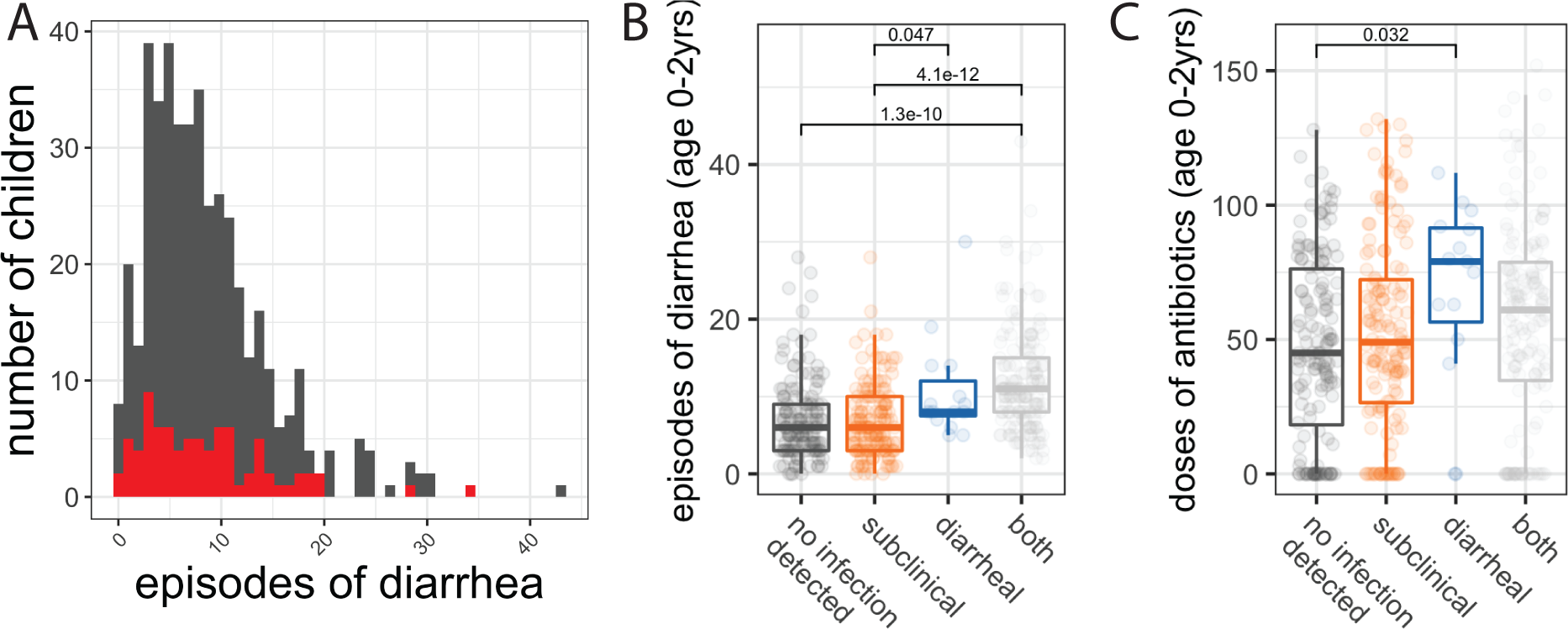
Diarrheal infection and antibiotic treatment were common and heterogenous in infants from Mirpur. **A**: Prevalence of diarrhea. Frequency of diarrheal episodes per child. Full Mirpur cohort in grey, red subset indicates the children whose samples were used in the microbiome study (and all subsequent figures). **B**: All-cause diarrhea was heterogenous among children with divergent *Cryptosporidium* outcomes. Number of diarrheal events per child based on cumulative *Cryptosporidium* status, both over the first two years of life. **C:** Antibiotic usage was heterogenous among children with divergent *Cryptosporidium* outcomes. Number of antibiotic events per child based on cumulative *Cryptosporidium* status, both over the first two years of life. Combination therapies were treated as separate doses. For B and C, the full cohort was used and statistics are shown if significant. For B and C, each box represents the median (inner line), 25^th^ percentile and 75^th^ percentile. Upper whiskers extend from the top of the box to the largest value within 1.5 times the interquartile range (distance between 25^th^ and 75^th^ percentile), and the lower whisker extends to the smallest value within 1.5 time the interquartile range. P-values were generated from a t.test without multiple testing correction.

Children who had at least one symptomatic episode of cryptosporidiosis had more cumulative episodes of diarrhea than children with exclusively subclinical infections or no *Cryptosporidium*-positive stool samples (**Figure 2B**). Additionally, children with only diarrheal episodes (i.e. no observed subclinical cryptosporidiosis) had more frequent exposure to antibiotics than children who had never tested positive for *Cryptosporidium* (**Figure 2C**). Frequent antibiotic use occurred (**Supplementary Figure 2A**), but there was no difference in antibiotic use during the month prior to infection between children with subclinical or diarrheal infections (**Supplementary Figure 2B**).

**Supplemental Figure 2:**
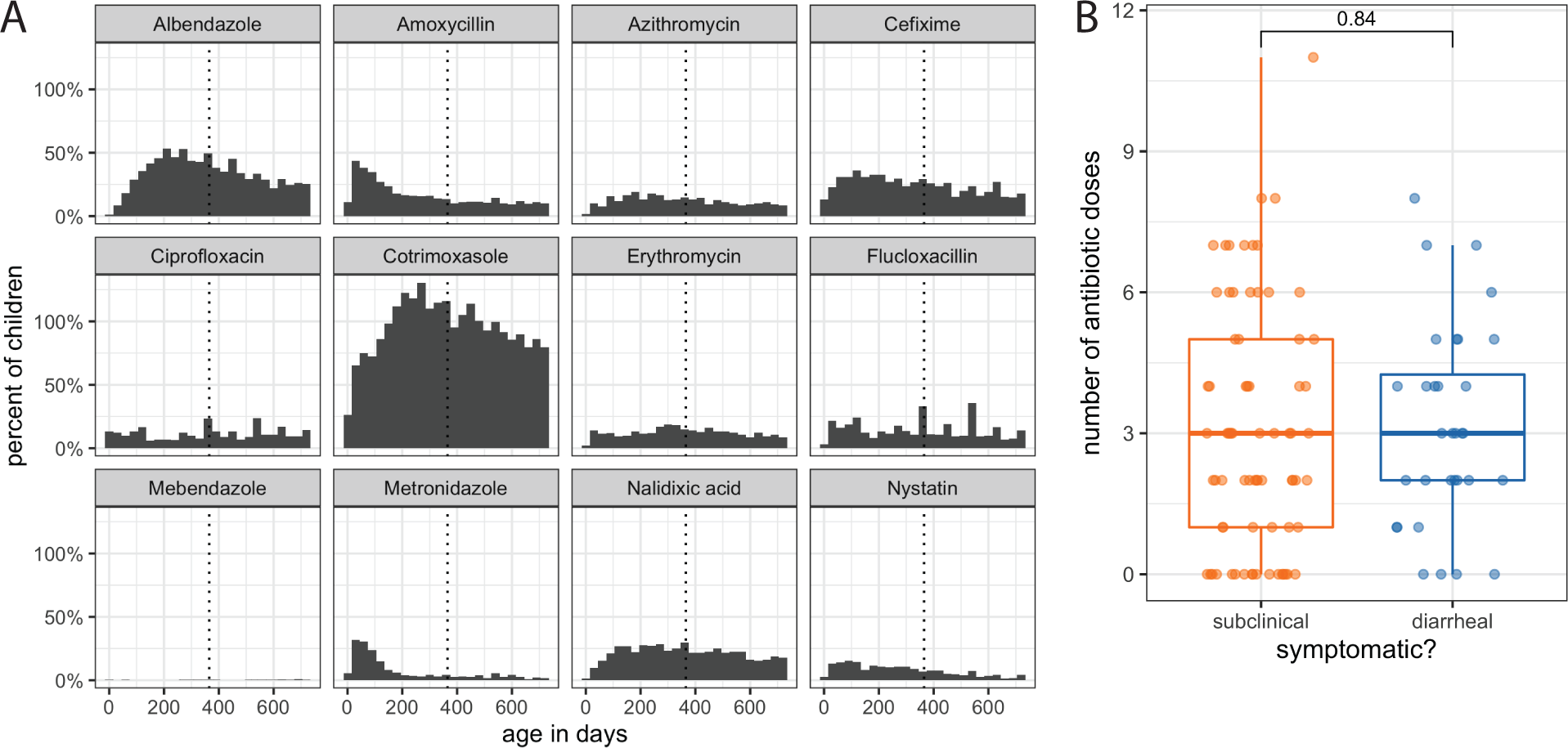
Frequent antibiotic use was age-dependent but homogenous between subclinical and diarrheal cases of *Cryptosporidium* in Mirpur. **A**: Antibiotic usage by drug. Antibiotics were prescribed for both diarrhea and other infections (e.g. respiratory). Combination therapies are listed for each subcomponent and multiple uses on a single drug within a month are listed as distinct episodes. Dotted line indicates the child’s first birthday. **B:** Children with diarrheal cryptosporidiosis in the microbiome sub-cohort took no more doses of antibiotics in the month leading up to the *Cryptosporidium*-positive stool sample than children with subclinical cryptosporidiosis, as determined using a t-test. Each box represents the median (inner line), 25^th^ percentile and 75^th^ percentile. Upper whiskers extend from the top of the box to the largest value within 1.5 times the interquartile range (distance between 25^th^ and 75^th^ percentile), and the lower whisker extends to the smallest value within 1.5 time the interquartile range.

### Microbiota Sequencing

Given the difference in all-cause diarrheal frequency between children with subclinical and diarrheal cryptosporidiosis (**Figure 2B**), we hypothesized that microbiome composition may influence the development of acute symptoms during cryptosporidiosis. 16S rRNA gene sequencing was performed on both the time-of-detection stool samples (*Cryptosporidium*-positive, including subclinical and diarrheal) and the corresponding surveillance stool collected immediately prior to the *Cryptosporidium*-positive sample (pre-detection; **Figure 1B**) for a subset of children who tested positive for *Cryptosporidium* (**Table 1**, red in **Figure 2A**). Pre-detection samples were collected within approximately one month of the time-of-detection samples (**Table 1**).

Sequencing produced 48,146,401 reads with a mean of 118,295.8 and median of 121,519 reads per sample (*raw reads* from **Supplemental Figure 1D**). Following quality filtration and taxonomy assignment, a mean of 27,809 reads per sample remained permitting us to subsample reads to a uniform depth of 10,000 reads per sample to correct for differences in sequencing depth across samples.

### Microbiota diversity

Following sequencing, taxonomy was assigned to reads using DADA2. Nearly 25% of reads were assigned to an amplicon sequence variant (ASV) belonging to the genus *Bifidobacterium* (**Figure 3A**) that represents a number of functionally-diverse species which colonize the infant gastrointestinal tract soon after birth. Microbiota alpha diversity measures (richness and evenness) were not statistically significantly different between sample groups (two-way ANOVA, post-hoc testing via Tukey’s Honestly Significant Difference method significance cutoff of p-value < 0.05; **Figure 3B-C**). Despite this lack of significance (*p* > 0.21 for all comparisons), the microbiota of infants who had diarrheal infection was, on average, less diverse than infants with subclinical infection, both prior to and at the time of infection (**Figure 3B-C**). Moreover, this cohort exhibited high inter-individual variation as many ASVs were specific to just a few children. Only a few ASVs were found in >50% samples (**Figure 3D**).

**Figure 3:**
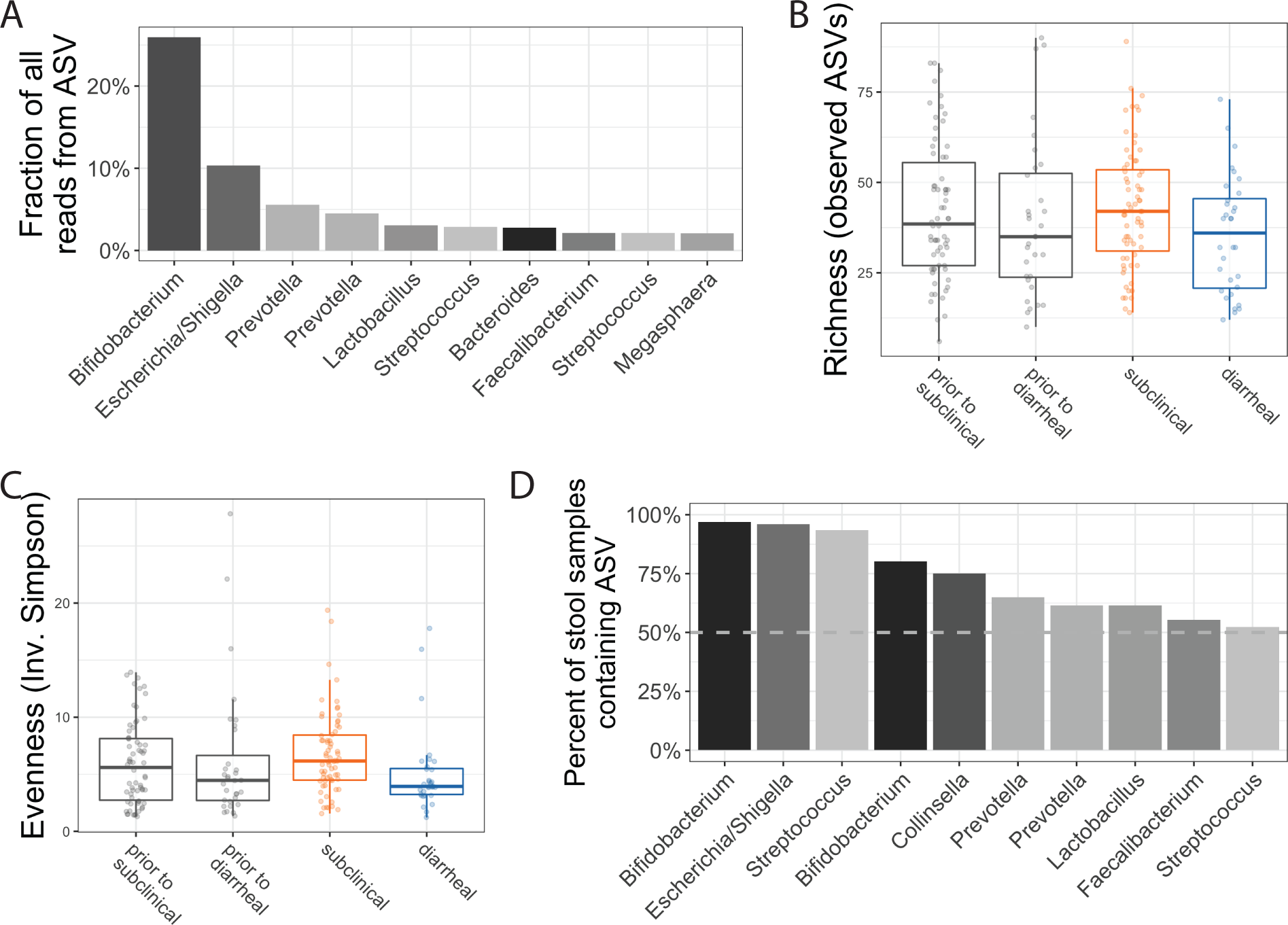
Microbiome samples were highly variable. **A**: Most abundant amplicon sequence variants (ASVs) in the study. Only the top 10 most abundant ASVs are shown. Nearly 25% of all reads were assigned to an ASV in the *Bifidobacterium* genus. **B**: Richness of each sample, or the number of ASVs present in a sample, was not significantly different across sample groups. For B and C, each box represents the median (inner line), 25^th^ percentile and 75^th^ percentile. Upper whiskers extend from the top of the box to the largest value within 1.5 times the interquartile range (distance between 25^th^ and 75^th^ percentile), and the lower whisker extends to the smallest value within 1.5 time the interquartile range. **C**: Evenness was also minimally different across sample groups. Evenness is a diversity metric calculated to represent how many different species are present and how well distributed those species are across samples; it is calculated using the inverse Simpson’s index. No significant differences in evenness was observed amongst any comparisons of clinical type (two-way ANOVA with multiple testing correction via Tukey’s Honestly Significant Difference method). **D**: Fraction of all samples containing a particular ASV, ordered by from highest to lowest. Very few ASVs were detected in many samples; however, almost all samples contain the most common *Bifidobacterium* ASV. ASVs: Amplicon sequence variants.

### Associations between diarrheal symptoms and the microbiota

To identify compositional differences in the microbiome among sample groups, principal coordinate analysis was performed using the Euclidean distance between samples. Pre-detection samples overlapped substantially with *Cryptosporidium*-positive samples and, among positive samples, subclinical and diarrheal samples did not separate (permutational multivariate analysis of variance using distance matrices [PERMANOVA], *p*>0.05, **Figure 4A**). The change in microbiota from pre-detection to time-of-detection for each child was similarly variable for both diarrheal and subclinical infections (PERMANOVA, *p*>0.05, **Supplemental Figure 3**).

Given the lack of separation between samples when considering overall microbiome composition, univariate analyses were used to identify individual ASVs that were significantly different between subclinical and diarrheal samples prior to and at the time of infection (**Figure 4B**). However, univariate statistics rely on assumptions of independence and, thus, may perform poorly with microbiome datasets due to correlations between and statistical interactions amongst members of the microbiota [52]. To make robust inferences of the importance of individual ASVs, we utilized a univariate approach designed specifically for sparse count data [53] as well as random forest classification to consider interactions amongst ASVs. Interpreting the results of these two approaches together provided a more stringent assessment of ASV importance.

Thus, classification using the random forest models was performed to determine if specific members of the microbiota were predictive of the development of diarrheal symptoms; important variables from the random forest models are highlighted on the volcano plots that also show the results of univariate statistical tests (**Figure 4B** & **C**). This machine learning approach was used to prioritize the results generated from univariate statistics. Classifier performance using the pre-detection or time-of-detection microbiome separately yielded predictive models (AUC > 0.6 for both prior to and at the time of infection **Figure 4C**); this performance was similar to the highest-performing classification models across a metanalysis of case-control clinical microbiome studies [54,55].

Both classifiers supported conclusions drawn by univariate analyses and identified several additional ASVs as important to classify subclinical and diarrheal samples (**Figure 4B** & **C**). Some important microbes for each classifier were not enriched in either sample group (large grey points in **Figure 4B**), this suggested that these ASVs are only important when analyzed in combination with others. Despite the effect of antibiotic treatment on the microbiota [56], the addition of a child’s antibiotic history did not significantly augment classifier performance (**Supplemental Figure 4**), indicating that there was no interaction between the important ASVs and antibiotic use. The infecting *Cryptosporidium* species (*C. hominis* or *C. meleagridis*) were not important variables in the random forest models, and child age was not an important variable in the time-of-detection model (**Figure 4D**).

**Supplemental Figure 4:**
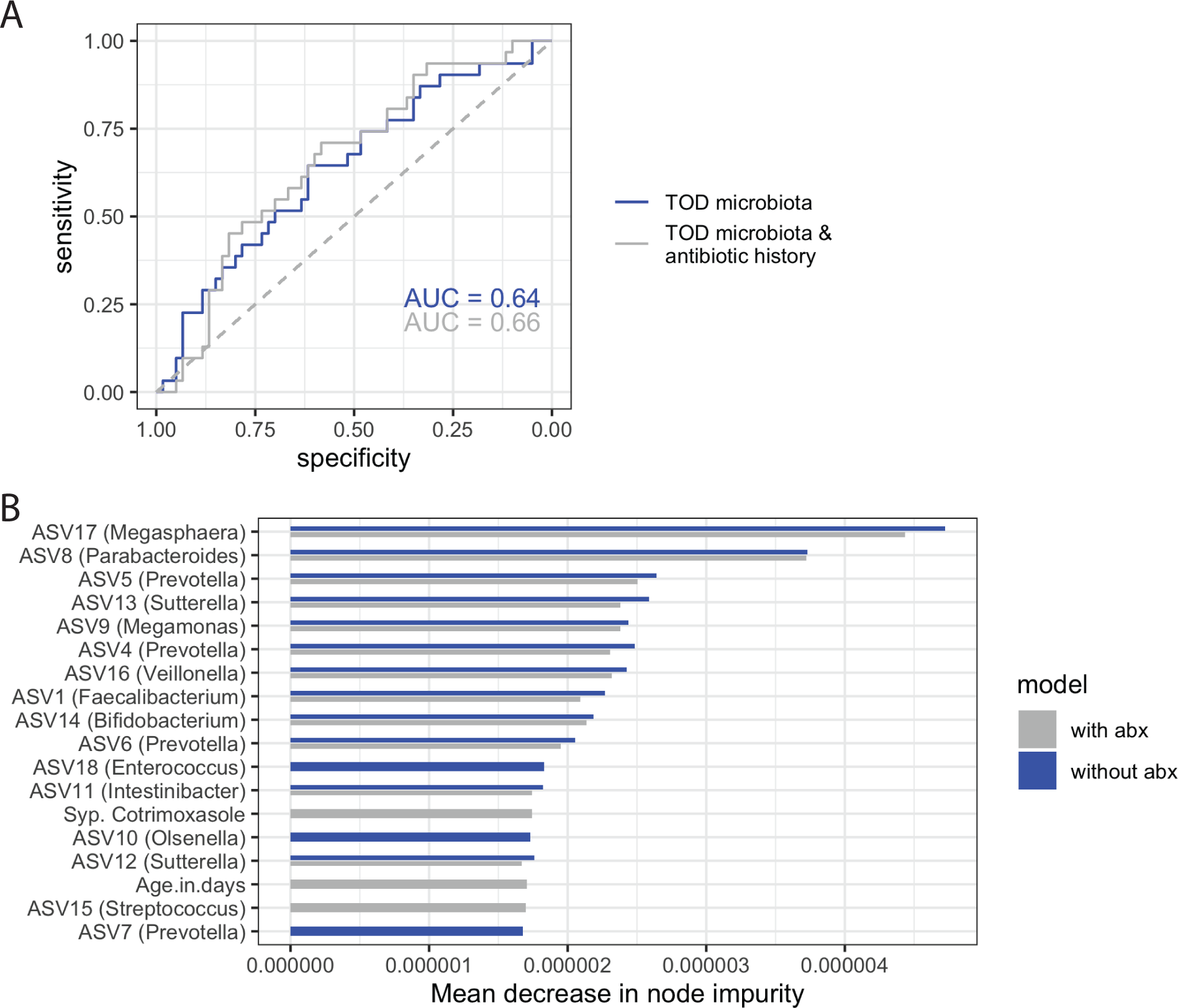
Difference in variable importance upon adding antibiotic history data. **A:** Classifiers built from the time-of-detection microbiota, with and without antibiotic history have similar performance (**A**) and identify the same important variables (**B**), indicating that antibiotic history did not add to the information gained from the microbiome.

We focused on ASVs that were identified via both the univariate statistics and machine learning approaches. For the pre-detection timepoint, these prioritized ASVs were assigned to the *Megasphaera, Flavonifractor, Morganella, Collinsella*, and *Lactobacillus* genera; for the time-of-detection timepoint, these included the same *Megasphaera* ASV, as well as ASVs assigned to *Parabacteroides, Enterococcus, Prevotella, Bifidobacterium, Sutterella, Veillonella, Megamonas*, and *Faecalibacterium* (**Figure 4B & D**). Combinations of ASVs were more predictive of diarrhea than any individual ASV, as evident by the similar Gini importance for all important variables (**Figure 3D**).

*Megasphaera*, in particular, was identified at both timepoints and both analytic approaches (**Figure 4B** & **E**). This *Megasphaera* ASV also accounted for at least 1% of reads across the entire study (**Figure 2A**). This bile acid-resistant species colonizes the small [57] and large intestines [58,59]. It can therefore be a major component of the microbiome at the site of *Cryptosporidium* parasite colonization. The other ASVs that contributed to model performance were either less abundant or resided predominantly in the large bowel. Although there were many environmental differences between the study sites, this ASV was also more likely to be detected at high abundance in our second study site, rural Mirzapur (**Figure 4F**), despite the observation that *Megasphaera* ASV did not vary with *Cryptosporidium* species (**Supplemental Material**). The most common *Cryptosporidium* species at Mirzapur was *C. meleagridis* rather than the *C. hominis* in Mirpur, but *C. meleagridis* has been associated with gastrointestinal disease in other studies and has also been shown to cause diarrhea in a human challenge experiment [60,61]. Children in Mirzapur were however less likely to develop diarrhea upon *Cryptosporidium* infection; 3% of *Cryptosporidium-*positive stools in Mirzapur were diarrheal, compared to 32% in Mirpur [1].

## DISCUSSION

Here, we identified differences in the microbiota composition and in the abundance of an individual ASV, *Megasphaera*, in infants who had either a subclinical or a diarrheal *Cryptosporidium* infection. Fecal samples from 72 *Cryptosporidium*-infected children in Mirpur, Bangladesh were used to profile the human microbiota during cryptosporidiosis (**Table 1, Figure 1**) with 16S rRNA gene sequencing (**Figure 3**). It is well established that the microbiome shifts with child development [62–64], and that it is highly variable in infants under the age of two [65– 67]. There was also universally frequent antibiotic use and enteric infections in this young population (**Figure 2C, Supplemental Figure 2, Table 1**). It was unsurprising therefore that there was a high degree of inter sample variability among these infants’ samples (**Figure 3A & D**).

Despite this variation, microbiome composition was predictive of diarrheal symptoms at the time of infection and up to a month prior (**Figure 4C**). Although individual members of the microbiome were associated with diarrhea (**Figure 4B**), no single ASV completely explained the clinical type of infection (**Figure 4D**). This observation is consistent with animal models of infection that have highlighted a complex relationship between the microbiota, host, and parasite [68–70]. For example, previous work found that antibiotics alone did not sensitize immunocompetent mice to infection [25], although certain probiotics [71], antibiotics [72], and deprivation of prebiotics [73] could exacerbate disease severity.

Higher abundance of one ASV, *Megasphaera* (class: Clostridia), was associated with subclinical *Cryptosporidium* infection whereas its absence or low abundance was more common in cases of *Cryptosporidium*-associated diarrhea (**Figure 4B & D**). *Megasphaera* was not associated with antibiotic use in this cohort (**Supplemental Figure 4**) or all-cause diarrhea (i.e. total number of diarrheal episodes, **Supplemental Material**). *Megasphaera* species can collocate in the small intestines [57] with *Cryptosporidium*, and were more frequently observed at high abundance in a community in which diarrhea was rarely seen during cryptosporidiosis (**Figure 4F;** [1]). *Megasphaera* are known to synthesize short chain fatty acids [74], compounds that regulate the intestinal homeostasis [75], impact the host immune response [76], and modulate osmotic diarrhea [77]. This ability of *Megasphaera* to produce short chain fatty acids or to modulate the host’s immune system through other mechanisms may be protective in attenuating disease outcome during *Cryptosporidium* infection. Alternatively, *Megasphaera* may be a biomarker for another microbiome- or immune-mediated mechanism of protection from diarrhea.

### Conclusion

In sum, the microbiome was predictive of *Cryptosporidium* diarrhea both prior to and at the time of infection. Low abundance of one member of the microbiome, *Megasphaera*, was associated with diarrheal symptoms. There is currently no effective drug for treating *Cryptosporidium* diarrhea in children and modulating members of the microbiota such as *Megasphaera* may be an appealing therapeutic strategy.

**SUPPLEMENTAL MATERIAL**, particularly analytic code, is available at https://github.com/maureencarey/cryptosporidium_microbiome.

## ACKNOWLEDGEMENTS

We thank the children and parents from the icddr,b study sites as well as the field workers, physicians, scientists, and staff of the Emerging Infectious Diseases Division of icddr,b for their key contributions to this research. We also wish to thank the members of the Petri lab group for their feedback, especially Jeff Donowitz, Kevin Steiner, Chelsea Marie, and Barbara Mann and the UVa Genome Analysis and Technology Core for conducting the sequencing for this paper.

## AUTHOR CONTRIBUTIONS

MAC and CAG conceived of and WP led the study. RH and ASGF founded the birth cohort and directed the Bangladesh studies. GLM and MAC conceived of the analyses. Field work and data collection at icddr,b were performed by MA and MK, with ASGF and RH providing supervision. DNA extraction and 16S rRNA library preparation were performed by MJU. UN oversaw the clinical database. GLM performed some preliminary microbiome sequence processing and machine learning analyses, with supervision by JP. MAC performed all final analyses. MAC drafted the manuscript. All authors edited and approved the final manuscript.

## DATA AVAILABILITY

Select clinical metadata is available on the NCBI’s dbGaP under accession number phs001665.v2.p1. The data for this study is collected as a sub-study of dbGaP phs001475.v2.p1. Raw sequence data generated for this study will be available in the Sequence Read Archive upon publication. Code for analysis is available at https://github.com/maureencarey/cryptosporidium_microbiome. Per the consent of the parents and guardians of the children in this study, all other deidentified data may be available upon request.

